# Ancient polyploidy and low rate of chromosome loss explain the high chromosome numbers of homosporous ferns

**DOI:** 10.1101/2024.09.23.614530

**Authors:** Zheng Li, Sylvia P. Kinosian, Shing H. Zhan, M. S. Barker

**Affiliations:** Department of Ecology and Evolutionary Biology, University of Arizona, Tucson, AZ 85721 USA; Big Data Institute, Li Ka Shing Centre for Health Information and Discovery, University of Oxford, OX3 7LF, United Kingdom

## Abstract

A longstanding question in plant evolution is why ferns have many more chromosomes than angiosperms. The leading hypothesis proposes that ferns have ancient polyploidy without chromosome loss or gene deletion to explain the high chromosome numbers of ferns. Here, we test this hypothesis by estimating ancient polyploidy frequency, chromosome evolution, protein evolution in meiosis genes, and patterns of gene retention in ferns. We found similar rates of paleopolyploidy in ferns and angiosperms from independent phylogenomic and chromosome number evolution analyses, but lower rates of chromosome loss in ferns. We found elevated evolutionary rates in meiosis genes in angiosperms, but not in ferns. Finally, we found some evidence of parallel and biased gene retention in ferns, but this was comparatively weak to patterns in angiosperms. This work provides genomic evidence supporting a decades-old hypothesis on fern genome evolution and provides a foundation for future work on plant genome structure.

## Introduction

A longstanding mystery of plant evolution is the origin of the exceptional chromosome number variation observed across the vascular plant phylogeny. Vascular plant chromosome numbers range from *n* = 2 in multiple angiosperms (Rice et al. 2015) to *n* = 1,260 in the eusporangiate fern, *Ophioglossum reticulatum* (Abraham and Ninan 1954). Reproductive mode is associated with variation in chromosome number across vascular plants. Homosporous ferns and lycophytes—plants that reproduce with one type of spore and develop bisexual gametophytes—are notable for their high chromosome numbers with an average gametic chromosome count of *n* = 57.05 (Klekowski and Baker 1966). In contrast, the angiosperms and heterosporous ferns which reproduce via mega- and microspores have an average haploid chromosome number of *n* = 15.99 (Grant 1963) and 13.62 (Klekowski and Baker 1966), respectively. Although a variety of cytogenetic mechanisms are known to influence chromosome number variation at lower taxonomic scales (Levin 2002; Mayrose and Lysak 2020), it is unclear what processes are responsible for the macroevolutionary pattern observed between homosporous and heterosporous plants.

Several hypotheses have been proposed to explain the origin and maintenance of high chromosome numbers in homosporous ferns (Wagner and Wagner 1980; Haufler and Soltis 1986; Barker and Wolf 2010; Manton 1950; Haufler 1987). The most compelling hypothesis invokes multiple rounds of whole genome duplication (WGD) and limit chromosome loss and limited gene deletion (Haufler and Soltis 1986; Haufler 1987). The alternatives, such as high ancestral chromosome numbers or ascending dysploidy are considered less likely because neopolyploid speciation is common in ferns (Otto and Whitton 2000; Wood et al. 2009) and cytological variation is predominantly euploid (Love, Love, and Pichi Sermolli 1977). An influential early hypothesis argued that most homosporous ferns were polyploids and that repeated cycles of genome doubling provided homoeologous heterozygosity, which was necessary to compensate for putatively high rates of inbreeding (Chapman, Klekowski, and Selander 1979; Klekowski and Baker 1966). However, isozyme studies demonstrated that homosporous ferns with the base chromosome number for their genus are diploid, not polyploid, with inheritance patterns and genetic variation consistent with outcrossing, not inbreeding (Haufler and Soltis 1986; Haufler 1987; Wolf, Haufler, and Sheffield 1987; Gastony and Darrow 1983; Gastony and Gottlieb 1982, 1985). To explain this intriguing combination of high chromosome numbers and diploid genetics, Haufler (1987) suggested that ferns experienced multiple rounds of polyploid speciation followed by gene silencing but not chromosome loss. In support of this hypothesis, a few studies have identified multiple silenced copies of nuclear genes in putatively diploid homosporous fern genomes (Pichersky, Soltis, and Soltis 1990; McGrath and Hickok 1999; McGrath, Hickok, and Pichersky 1994), and the active process of gene silencing without chromosome loss in a polyploid genome (Gastony 1991). However, linkage mapping of the diploid homosporous fern, *Ceratopteris richardii* (*n* = 39) failed to identify homoeologous chromosomes (Nakazato et al. 2006).

Despite the lack of clear evidence for paleopolyploidy in linkage maps of *Ceratopteris* (Nakazato et al. 2006), recent genomic analyses found evidence for ancient genome duplications in ferns (Barker 2012; Vanneste et al. 2015; F.-W. Li et al. 2018; One Thousand Plant Transcriptomes Initiative 2019; Clark, Puttick, and Donoghue 2019; Barker 2009; Pelosi et al. 2022; Chen et al. 2022; Marchant et al. 2022; Fang et al. 2022; Huang et al. 2022). Thus, it is no longer a question that ferns experienced ancient polyploidy, but whether the frequency of ancient WGDs in ferns was significantly higher than other lineages of vascular plants and whether that could explain high chromosome numbers in ferns.

Here, we explore how cycles of polyploidy have interacted with rates of chromosomal evolution to produce the high numbers of chromosomes observed in homosporous ferns. By incorporating inferences of paleopolyploidy from previous studies (One Thousand Plant Transcriptomes Initiative 2019; Pelosi et al. 2022; Huang et al. 2020; Chen et al. 2022; Shen et al. 2018; F.-W. Li et al. 2018; Marchant et al. 2022; Huang et al. 2022), we compared the incidence of WGD in ferns to other lineages of vascular plants. We then evaluated differences in the rates of chromosomal evolution among these clades by estimating the modes and rates of chromosome number evolution among more than 1,900 vascular plant genera. Using these analyses, we test whether ferns have 1) an extraordinary amount of WGD in their history with similar rates of chromosomal evolution as other lineages, 2) a similar rate of WGDs but a relatively slow rate of dysploidy compared to other plants, or if 3) ferns have ancestrally high chromosome numbers and a relatively low number of rounds of polyploidy. Finally, considering the large numbers of chromosomes involved in fern meiosis, we tested whether the rates of protein evolution for meiosis-related genes are significantly different between ferns and angiosperms. Together, our inferences provide complementary views on the evolution of high chromosome numbers in homosporous ferns.

## Results

### Inferences and frequency of paleopolyploidy

To infer ancient WGDs in ferns, we assembled a transcriptomic data set of 133 fern species from published studies (Fig. 1 and Supplementary Table 1). Using gene age distributions, we found evidence for peaks of gene duplication consistent with WGDs in all fern transcriptomes (Supplementary Table 1). We also used ortholog divergence analyses to place some ancient WGDs in ferns in a phylogenetic context. These WGDs were not analyzed by the One Thousand Plant Transcriptomes Initiative (Supplementary Table 2). To explore the frequency of paleopolyploidy in ferns, we incorporated inferences and phylogenetic placements of ancient WGDs from this study and other published studies (Pelosi et al. 2022; Huang et al. 2020; Chen et al. 2022) (Supplementary Table 3). As tree topology is known to vary between studies with different markers (e.g., Hasebe et al. 1995; Schneider et al. 2004; Testo and Sundue 2016; PPG I 2016; McKibben, Finch, and Barker 2024), the placement of WGDs differed slightly across the studies used here. We used the K_s_ values to infer which WGDs matched across studies. If a *K*_s_ value was within 0.1 across all studies examined, we considered it to be the same WGD (see Fig. 1). One study (One Thousand Plant Transcriptomes Initiative 2019) inferred a WGD shared by all ferns at the base of the monilophyte phylogeny (WGD 1, see Supplementary Table 3). However, other studies have not inferred this WGD and there is no syntenic evidence for this putative ancient WGD. In order to remain conservative in our estimates, we did not count this in our total estimate of WGDs for ferns. Overall, we found evidence for 30 putative ancient WGDs throughout the evolutionary history of monilophytes (Fig. 1 and Supplementary Fig. 1).

**Fig. 1.**
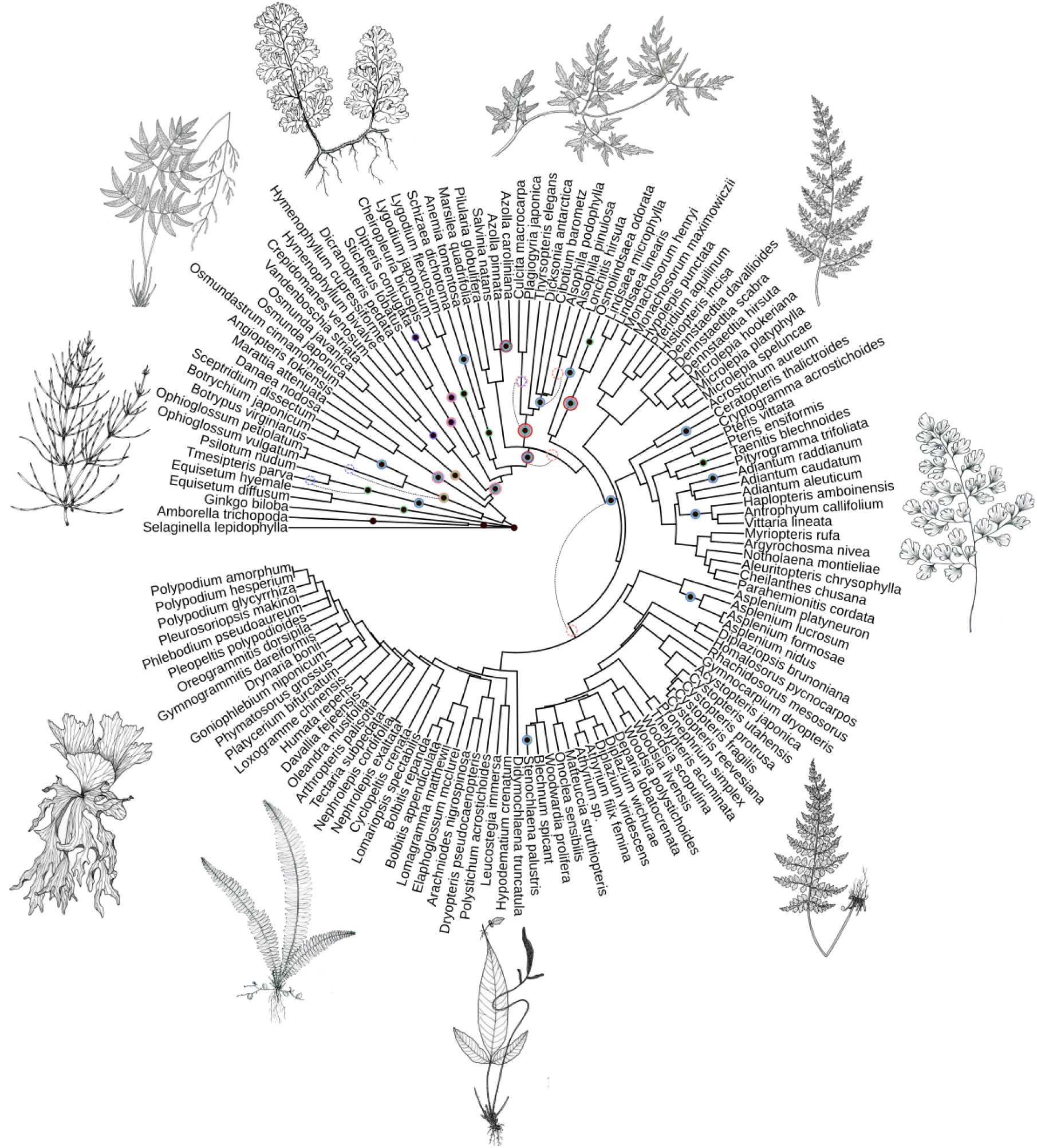
Phylogenetic placement of ancient WGDs inferred in the phylogeny of monilophytes. Black dots indicate whole genome duplications inferred for pteridophytes. The black dots at the base of the tree show WGD in outgroups. Circles around the dots show support for a duplication from different studies as follows: green, analyses from this study using 1KP data; blue, Pelosi et al. 2022; pink, Huang et al. 2020; red, Chen et al. 2022. Dotted circles show where previous authors placed WGD in their trees, but the Ks values of those WGD are very close to what we inferred, so we believe they are the same duplication (dotted lines connect the two). This phylogeny was modified from Testo and Sundue, 2016. Illustrations of ferns are credited to Yifan Li.

We estimated the frequency of paleopolyploidy by calculating the mean of the number of inferred ancient WGDs in the ancestry of each species (Fig. 2a). On average, fern species experienced 2.81 ± 0.75 rounds of ancient genome duplication. Similar to ferns, the genome of an angiosperm species went through an average of 3.5–3.9 rounds of ancient WGD (One Thousand Plant Transcriptomes Initiative 2019; McKibben, Finch, and Barker 2024). Gymnosperm and lycophyte species experienced lower numbers of ancient WGDs, with averages of 1.86 ± 0.34 and 1.52 ± 1.21 rounds of ancient WGD, respectively (Fig. 2a; One Thousand Plant Transcriptomes Initiative 2019). Interestingly, heterosporous ferns experienced 2.4 ± 0.54 rounds of ancient WGD whereas homosporous ferns experienced 2.82 ± 0.75 rounds of ancient WGD. It is important to note that the sample sizes here were very different, with only 5 heterosporous taxa compared to 128 homosporous taxa.

**Fig. 2.**
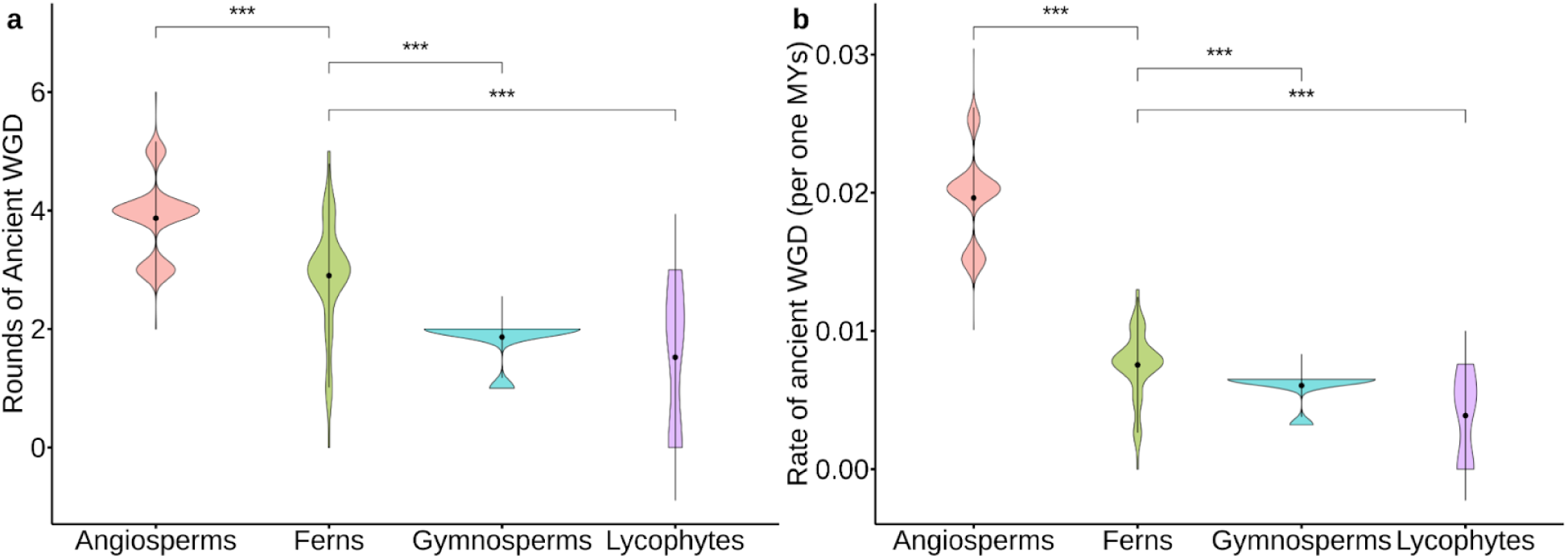
The number of inferred ancient WGDs in vascular plants. **a**, The number of inferred ancient polyploidization events within each lineage is shown in the violin plots. The sample sizes for angiosperms, ferns, gymnosperms, and lycophytes are 674, 133, 81, and 21, respectively. Inferred ancient polyploidization events in angiosperms, gymnosperms, and lycophytes are based on estimates from 1KP. **b**, The rate of inferred ancient polyploidization events within each lineage is shown in the violin plots. The rate of inferred WGD is estimated by the number of inferred WGD divided by the minimum crown group age of each lineage. The minimum crown group age used for angiosperms, ferns, gymnosperms, and lycophytes are 197.5, 384.9, 308.4, and 392.8 million years, respectively. The black dot indicates the mean, the thick black bars represent the standard deviation of the data, the color shading represents the density of data points. The comparisons of the mean of ferns to other lineages by the two-sample Mann-Whitney U test are provided. ‘***’ represents *p* < 0.0001.

We found a significant difference in the number of ancient WGDs in the ancestry of fern and angiosperm species (U = 14,011.5, *p* = 0; Mann-Whitney U test) (Fig. 2a). The number of rounds of ancient WGD in ferns was also significantly different compared to gymnosperms (U = 9,307, *p* < 10^−15^) and lycophytes (U = 2,262, *p* < 10^−6^) (Fig. 2a). To account for the impacts of age on the number of ancient WGDs in each lineage, we also estimated the rate of ancient WGD/Ma using the number of ancient genome duplications divided by the minimum crown group age (Fig. 2b). The minimum crown group age used for angiosperms, ferns, gymnosperms, and lycophytes were 197.5, 384.9, 308.4, and 392.8 Ma, respectively (Supplementary table 4) (Morris et al. 2018). We estimated the rates of ancient WGD at 0.0073 ± 0.002 WGD/Ma in ferns and 0.0196 ± 0.003 WGD/Ma in angiosperms. In gymnosperms and lycophytes, the rates were 0.0060 ± 0.001 and 0.0039 ± 0.003 WGD/Ma, respectively (Fig. 2b). We found the rate of ancient WGD in flowering plants was more than twice that of the ferns (U = 89,597, *p* = 0). In contrast, the rate of WGD in the gymnosperms and lycophytes was significantly lower than the ferns (U = 8,681, *p* < 10^−14^; U = 236, *p* < 10^−11^, respectively) (Fig. 2b).

### Patterns and rates of chromosome number evolution

Using ChromEvol, we explored chromosome number evolution across *rbcL* phylogenies of 1,918 angiosperm, 183 fern, and 50 gymnosperm genera for which chromosome numbers were available. We omitted lycophytes in this analysis because of the small number of genera; however, we used them as an outgroup for the phylogeny. Our ChromEvol rates (from the best models) were similar to our estimates in the genomic analyses, with rates of ancient WGD of 0.0096 WGD/Ma in ferns, 0.0129 WGD/Ma in angiosperms, and 0.0010 WGD/Ma in gymnosperms (Fig. 3, Supplementary table 5). As with our estimates from genomic inferences, angiosperms were found to have 1.4 times the rate of polyploidy as ferns and 13.05 times the rate of polyploidy found among gymnosperms (Fig. 3, Supplementary table 5). Angiosperms were also estimated to have higher rates of dysploidy than ferns and gymnosperms, with both descending dysploidy and ascending dysploidy occurring more frequently in angiosperms than ferns and gymnosperms. Overall, the estimated rates for all types of chromosome number evolution in ferns and gymnosperms were lower than in angiosperms, and gymnosperms were the lowest among the vascular plants.

**Fig. 3.**
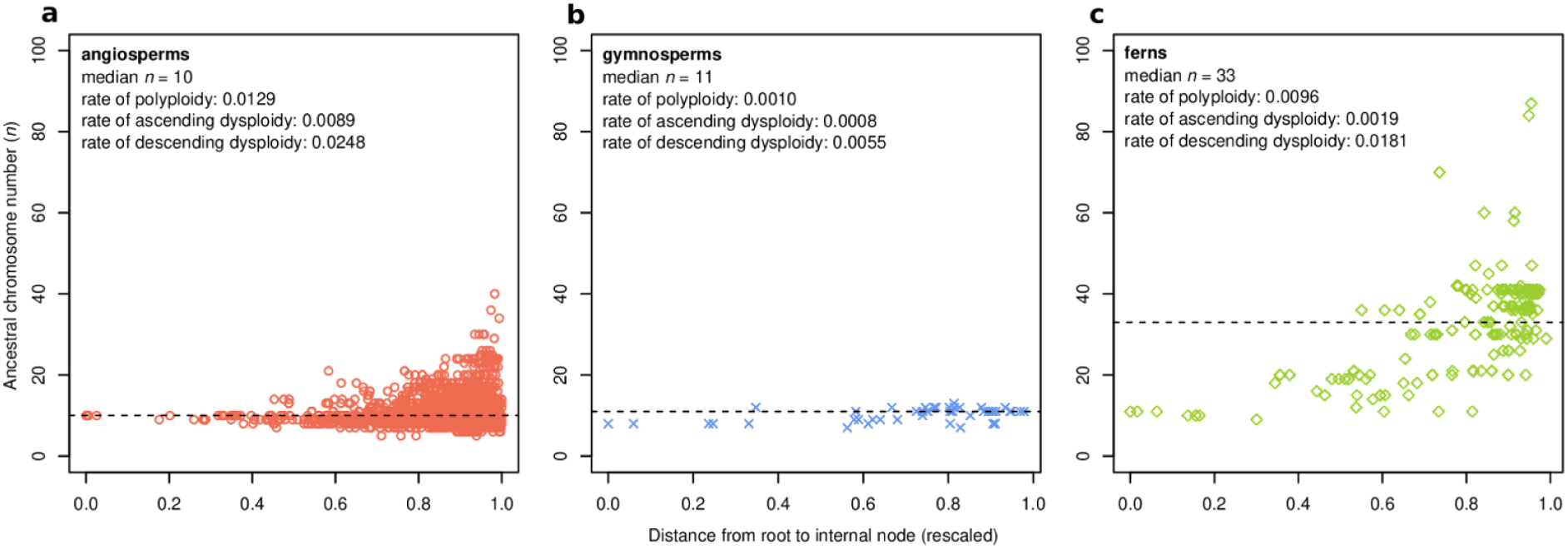
Patterns of inferred ancestral chromosome number in angiosperms, gymnosperms, and ferns. The x-axis represents the root-to-internal node distance on the phylogenies analyzed using the best model supported by ChromEvol (Supplementary Table 5). The y-axis shows the ancestral chromosome numbers taken from the ChromEvol results. The dotted line shows the median chromosome number for each group. The rate of polyploidy, ascending and descending dysploidy are presented in the figure legends. The pattern in (a) angiosperms, (b) gymnosperms, and (c) ferns is shown by 1,918 data points, 51 data points and 199 data points, respectively.

To further explore the pattern of chromosome number evolution among seed plants and ferns, we plotted the inferred ancestral chromosome numbers at each internal node of the *rbcL* phylogenies in our phylogenetic reconstruction of chromosome evolution (Fig. 3). Given current tools it is difficult to quantify uncertainty for ChromEvol estimate, but we plotted the posterior complement as a metric of uncertainty (Supplementary Fig. 6). Our reconstructions yielded three different patterns of chromosome number in these lineages. In angiosperms, chromosome numbers appear to be largely maintained over their evolutionary history, with inferred numbers at most nodes near the base chromosome number for the clade. This pattern is recovered even near the tips of the angiosperm phylogeny where signatures of recent polyploidy events are most likely to be observed. These estimates suggest that most angiosperms quickly reduce chromosome number following polyploidy (Fig. 3a). In contrast, gymnosperms demonstrated almost no variation in inferred chromosome number over their phylogeny. Nearly all nodes in the gymnosperm phylogeny had an inferred chromosome number of *n* = 11 or 12 with evidence of recent chromosome loss (Fig. 3b). The chromosome number of ferns increased towards the tips of the phylogeny. In particular, at least two distinct “steps” of chromosome number increase were evident in our analysis (Fig. 3c). Although our inferred fern chromosome numbers indicated some dysploidy, there appears to be remarkable conservation of chromosome numbers among many nodes following these broadly shared increases in chromosome number (Fig. 3c).

### Rates of protein evolution of meiosis genes in angiosperms and ferns

High chromosome numbers in polyploids can lead to multivalent formation and failure in chromosome pairing during meiosis (Comai 2005; Bray et al. 2024; Bomblies et al. 2016). It is not known how homosporous ferns overcome these potential challenges in meiotic pairing and stability. Previous studies have found evidence for the adaptive evolution of genes responsible for meiotic stabilization in autotetraploid *Arabidopsis arenosa* (Yant et al. 2013; Morgan et al. 2020), and kinetochore components in autotetraploid *Cochlearia* (Bray et al. 2024). A recent study in *Solanum* also showed elevated rates of protein evolution in reproductive proteins (Moyle, Wu, and Gibson 2021). Considering the challenges of meiosis with such high chromosome numbers in ferns, we hypothesized that ferns may have adapted to larger numbers of chromosomes and we may see increased rates of protein evolution of meiosis-related genes. Leveraging the phylogenomic data available, we tested if the rates of putative meiosis-related genes in ferns were elevated compared to a randomly selected pool of non-meiosis genes. We also conducted a similar analysis in angiosperms as an indirect comparison to ferns.

To build a gene list of putative meiosis-related genes in ferns, we first identified 352 meiosis-associated genes based on the *Arabidopsis* Gene Ontologies (GO) category (GO:0007126). We also randomly selected 500 non-meiosis genes in *Arabidopsis* as a background for comparison. Genes from both lists were used as the query to BLASTP with an E-value of 1e-20 as cutoff against eleven angiosperm genomes and ten fern genomes and transcriptomes as databases. The *Physcomitrella patens* genome was used as an outgroup. Gene families were clustered with Orthofinder 2.3.7 (Emms and Kelly 2015) using the homologous sequences from the initial BLASTP research. We then filtered gene families with the presence of at least one *P. patens* sequence and one *A. thaliana* sequence to build gene family phylogenies. Overall, 129 and 720 gene family trees were constructed for meiosis-related genes and randomly selected background genes, respectively.

To account for different rates of molecular evolution between angiosperms and ferns, we compared the rate of protein evolution for meiosis-related genes to the background set of genes within each lineage. After re-rooting with *Physcomitrella patens* and dropping tips of the outgroup, we estimated the mean, minimum, and maximum root-to-tip distance for each species in each gene tree. We used the Mann-Whitney *U* test to compare the distribution of root-to-tip distance between meiosis and background genes in angiosperms. We then performed the same test between meiosis and background genes in ferns. In angiosperms, we observe a significantly higher rate of protein evolution in meiosis genes compared to the background genes using maximum root-to-tip distance (U = 2,363,289.0, *p* < 0.001) (Fig. 4, Supplementary table 6). However, we found no significant difference in the rate of protein evolution in meiosis-related genes compared to the background genes in ferns (U = 2,113,199.5, *p* = 0.1710). This overall pattern was consistent when using the mean, minimum, and maximum root-to-tip distance, and with or without outliers from the distribution (Fig. 4, Supplementary table 6).

**Fig. 4.**
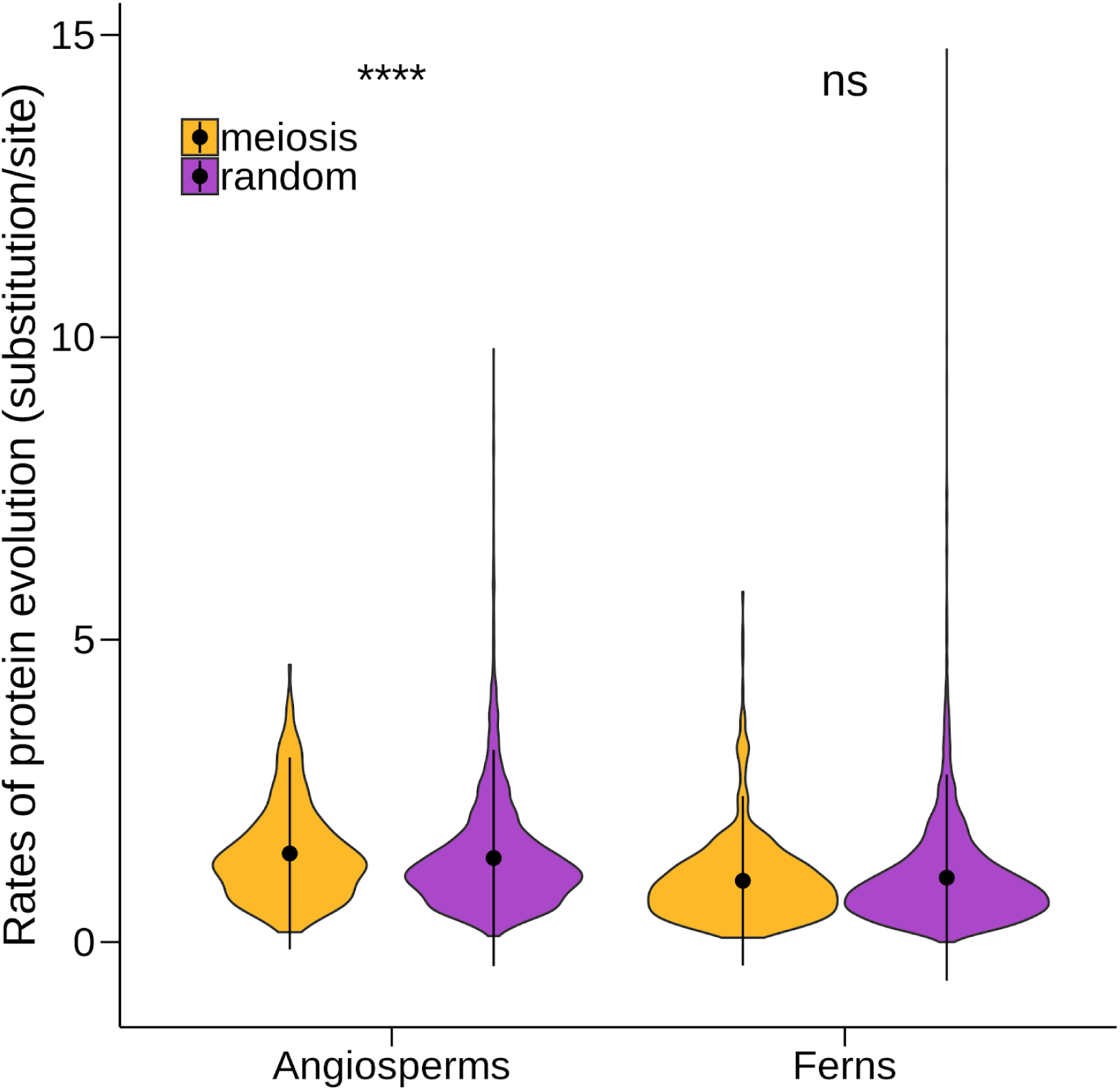
The rates of protein evolution of meiosis-related genes in angiosperms and ferns. The rates of protein evolution of meiosis-related genes and randomly selected genes of angiosperm and ferns are shown in the violin plots. The black dot indicates the mean, the thick black bars represent the standard deviation of the data, and the color shading represents the density of data points. In total, 129 and 720 gene family phylogenies were constructed for meiosis-related genes and randomly selected genes in angiosperms and ferns. For each gene tree, we estimate the maximum root-to-tip distance for each species. The comparisons of the mean of meiosis and random genes by the two-sample Mann-Whitney U test are provided. ‘****’ represents *p* < 0.0001.

### Biased gene retention and loss following inferred WGDs

Biased gene retention and loss is a common feature of ancient WGDs (Li et al. 2020), but limited studies have investigated this in monilophytes (Pelosi et al. 2022). To test for biased gene retention and loss among the inferred fern paleopolyploidies, we compared the overall differences between the GO composition of retained paralogs and whole transcriptomes using a principal component analysis (PCA) on the number of genes annotated to each GO category. The PCA found two significantly different clusters (*p* < 0.001) with some overlap (Fig. 5). The retained paralogs formed a tighter cluster with a narrower 95% confidence interval compared to the whole transcriptome (Fig. 5). We also used a hierarchical clustering approach to assess the overall GO composition similarity of paralogs retained from ancient WGDs. We observed biased retention and loss from all inferred WGDs in monilophytes (Supplementary Fig. 2-3). However, we found little evidence of parallel patterns of gene retention across different ancient genome duplications, similar to the findings of Pelosi et al. (2022). Most paleopolyploidy events were not resolved based on the hierarchical clustering approach (Supplementary Fig. 2). We further used a simulated chi-square test to infer whether any categories were significantly over- or under-retained. Many WGDs in ferns were significantly enriched for genes associated with transcription factor activity, DNA or RNA binding, and the nucleus (Supplementary Fig. 3). Paralogs retained from these ancient duplications were often significantly under-retained for genes associated with other enzyme activity, hydrolase activity, and transferase activity (Supplementary Fig. 2-3). Overall, all ancient WGDs in ferns had a pattern of biased gene retention and loss, but parallel patterns of retention and loss as predicted by the Dosage Balance Hypothesis (Papp, Pál, and Hurst 2003) were observed in only a few ancient genome duplications.

**Fig. 5.**
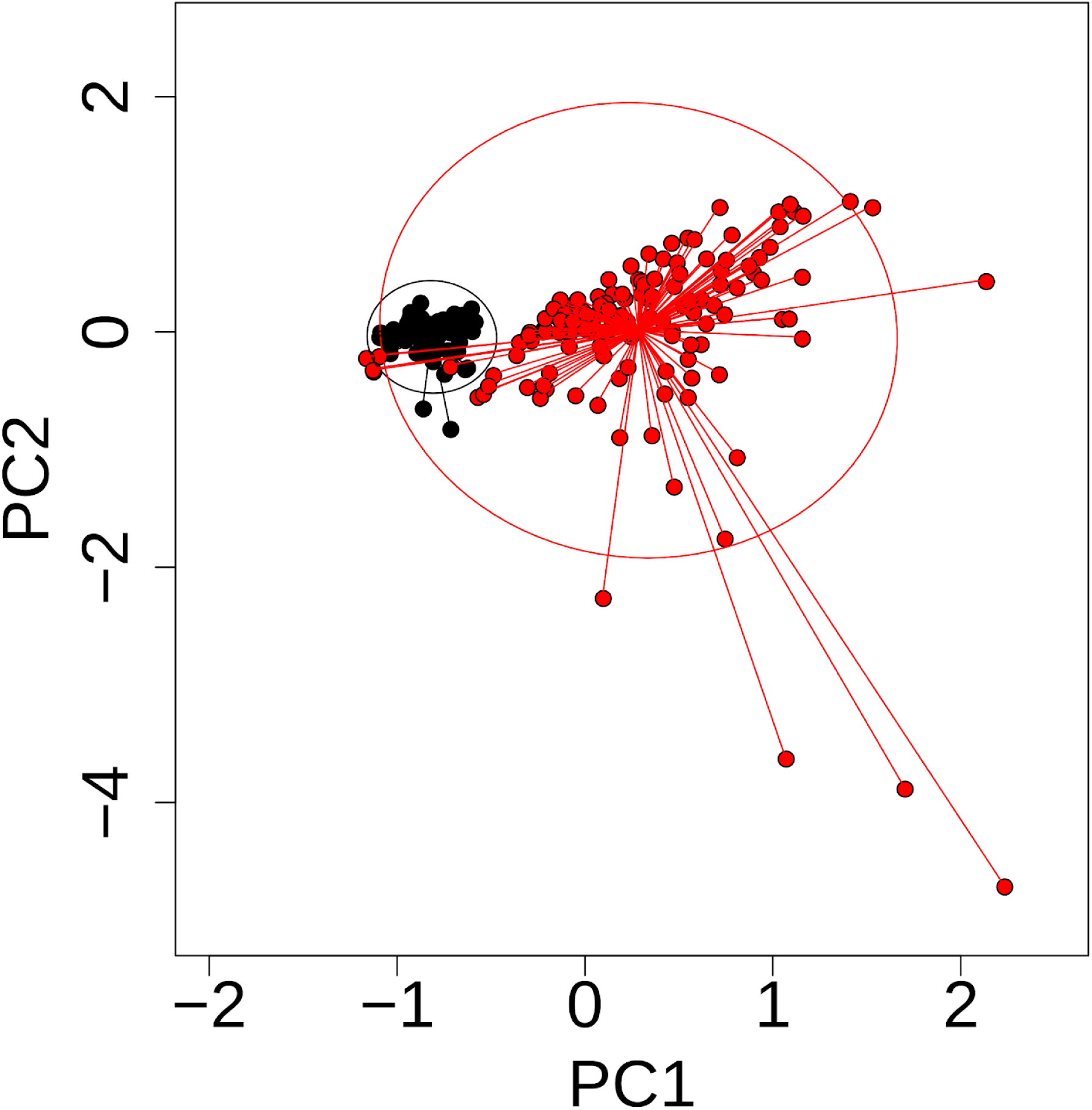
Principal component analysis of the GO category composition of all genes in each genome/transcriptome and WGD paralogs. Red circles indicate the number of genes annotated to each GO category in the whole transcriptomes; black circles, number of WGD paralogs annotated to each GO category. Ellipses represent the 95% confidence interval of SD of point scores.

## Discussion

Our genomic and phylogenetic analyses provide new tests of long-standing hypotheses of fern genome evolution. Across four studies (Huang et al. 2020; Chen et al. 2022; Pelosi et al. 2022; One Thousand Plant Transcriptomes Initiative 2019) and new analyses, we found evidence for 30 putative ancient WGDs throughout the monilophyte phylogeny (Fig. 1). The phylogenetic placement of these inferred WGDs indicates that ferns, on average, experienced 2.81 rounds of ancient polyploidy in their ancestry or 0.0073 WGD/Ma. Strikingly, our phylogenetic reconstruction of chromosome number evolution yielded a similar average rate of ancient polyploidy in ferns—0.0096 WGD/Ma—with completely different data. The quantitatively similar estimates from the two different inference methods and datasets suggest these results are robust and complementary. Our genomic and chromosomal phylogenetic analyses also found that angiosperms have the highest rates of ancient WGD and dysploidy among vascular plants. However, the average number of ancient WGDs per species is similar in ferns and angiosperms: 2.81 compared to ~3.7 (One Thousand Plant Transcriptomes Initiative 2019; McKibben, Finch, and Barker 2024). Further, we estimated the average rate of chromosome loss in ferns has been about half the rate for angiosperms across their phylogenies. This lower estimated rate of chromosome loss among ferns is consistent with their much higher number of chromosomes compared to angiosperms (Klekowski and Baker 1966; Carta, Bedini, and Peruzzi 2020; Kinosian, Rowe, and Wolf 2022; Nakazato et al. 2008). Taken together, our different inferences support the hypothesis that the high chromosome numbers of homosporous ferns are a product of multiple rounds of ancient polyploidy coupled with a relatively slow rate of chromosome loss (Haufler 1987; Gastony 1991; Haufler and Soltis 1986; Barker 2013).

Alternative hypotheses have been proposed to explain the high chromosome numbers of homosporous ferns that do not involve polyploidy but instead attribute high numbers as the ancestral state (Duncan and Smith 1978) or the result of ascending chromosomal fission (Wagner and Wagner 1980; Haufler and Soltis 1986; Barker and Wolf 2010). However, these explanations have not gained much support given the high frequency of recent polyploidy in ferns (Wood et al. 2009) and the uniform size and structure of fern chromosomes across genera (Wagner and Wagner 1980; Liu et al. 2019). In our study, we observed increases in chromosome number towards the tips of the fern phylogeny (Fig. 3). This result indicates that numbers have increased over time and rejects the hypothesis of ancestrally high chromosome numbers in ferns. The rate of ascending dysploidy was an order of magnitude lower than descending dysploidy in ferns, and both were lower than the rates of dysploidy in angiosperms. These results reject the second alternative hypothesis of ascending chromosomal fission. Considering the frequency of recent and ancient polyploidy among ferns and their relatively slow rates of dysploidy, neither of these alternative hypotheses is well supported by our results.

Our results do support the hypothesis that reproductive mode may be responsible for chromosome number disparity in ferns (Kinosian and Barker 2024; Kinosian, Rowe, and Wolf 2022). It is well-established that homosporous and heterosporous ferns have significantly different chromosome numbers (e.g., Klekowski and Baker 1966), and here we show that they have a similar number of rounds of WGD, with 2.82 in homosporous ferns and 2.4 in heterosporous ferns. This is consistent with our contention that rapid chromosome loss in heterosporous (pteridophyte or angiosperm) but not homosporous lineages of vascular plants is responsible for the differences in chromosome numbers among these lineages. A potential mechanism for this disparity in chromosome number between heterosporous and homosporous lineages could be female meiotic drive, which would only be able to act in the asymmetrical female meiosis of angiosperms and heterosporous ferns (Pardo-Manuel de Villena and Sapienza 2001). Specially, female meiotic drive can cause rapid evolution of centromere size and structure (Talbert and Henikoff 2022), karyotype (Blackmon et al. 2019), and chromosome number (Burt and Trivers 2009; Fishman et al. 2014; Plačková et al. 2024)Meiosis is symmetrical in homosporous ferns, eliminating the possibility for female meiotic drive and any subsequent genomic changes (Kinosian and Barker 2024). Pteridophytes are a perfect natural study system for investigating these dynamics as there are sister clades with heterosporous and homosporous species in both ferns and lycophytes.

Our analysis of evolutionary rates of meiosis-related genes revealed no evidence that fern meiosis has broadly adapted to accommodate high numbers of chromosomes. We found that meiosis-related genes in flowering plants generally evolved faster compared to other genes in the genome, but this was not the case for meiosis-related genes in ferns (Fig. 4). These results are consistent with recent research that has found evidence for meiotic adaptation to polyploidy among different flowering plant species (e.g, Hollister 2015; Yant et al. 2013; Bomblies 2023; Bray et al. 2024; Morgan et al. 2020). Interestingly, there is some evidence of gene family expansion within angiosperm and heterosporous pteridophyte lineages (Dhakal, Wolf, and Harkess 2024), but it is unclear the function of these particular gene families. Considering that ferns and flowering plants have similar histories of polyploidy, some of the differences in chromosomal and genome evolution between angiosperms and ferns may be caused by the evolution of meiosis-related genes. It may be that novel meiotic adaptations in angiosperms restore bivalent pairing or stabilize the genome following polyploidy (e.g., (Bomblies and Madlung 2014; Gonzalo 2022) ultimately result in chromosome number reductions during flowering plant diploidization. In contrast, fern meiosis proteins appear to be more broadly conserved than their angiosperm homologs and this conservation could play a role in the slower rate of chromosome loss during homosporous fern diploidization. Notably, the patterns of meiotic gene rate evolution are consistent with meiotic drive; faster meiotic protein evolution in heterosporous lineages with meiotic drive but conserved meiotic proteins in homosporous lineages without meiotic drive (Kinosian and Barker 2024; Zedec et al 2016). Whether or not the broad patterns of post-polyploid genome evolution in flowering plants and ferns result from the differences we observe in the rates of meiotic gene evolution is unclear from our present analyses. Our results do suggest that there are different selection pressures on genes associated with meiosis in ferns and angiosperms. Future work is needed to better understand the fundamental differences between meiosis and meiosis-related gene evolution among lineages of vascular plants, and consider the possible mechanisms causing these differences.

To further investigate the unique genome evolution in ferns, we evaluated the pattern of gene retention and loss following ancient polyploids. Previous angiosperm studies have found parallel gene retention in single ancient WGDs (Scannell et al. 2007; Mandáková et al. 2017), and multiple independent ancient WGDs (Schranz and Mitchell-Olds 2006; Barker et al. 2008; Mandáková et al. 2017; Z. Li et al. 2018). Fewer studies have investigated biased gene retention in ferns (Zhang et al. 2019; Pelosi et al. 2022). Recent work by Pelosi et al. (2022) found biased gene retention of GO categories using fern transcriptomes. In the present study, we found similar patterns of gene retention and loss in ferns using GO composition analyses and a combination of whole genomes and transcriptomes (Fig. 5). This parallel gene retention found by our study and Pelosi et al. (2022) is not as dramatic as previously observed in flowering plants using a similar statistical framework (Barker et al. 2008; Mandakova et al. 2017). This result may be expected for two reasons. First, the phylogenetic depth of the fern phylogeny might weaken the parallel retention pattern. Extant fern orders share a most recent common ancestor that is estimated to be ~350 MYA (Rothfels et al. 2015; Morris et al. 2018). Given that each fern species has experienced 2.81 rounds of ancient WGDs on average, many signals of gene retention might also be imbricated by different types of duplications (Pelosi et al. 2022). Second, a lower chromosome loss rate and lineage specific diploidization in ferns might result in weaker parallel retention compared to angiosperms. Lower rates may drive more lineage specific gene losses in these lineages as gene losses accumulate more slowly and idiosyncratically across the phylogeny rather than rapidly in a common ancestor. Consistent with this pattern, we recently found that the rate of gene loss and fractionation after WGD is not correlated with WGD age in ferns and lycophytes, but it is among angiosperms (Z. Li et al. 2020). The strong parallel retention pattern in angiosperms might be driven by their high rates of chromosome evolution. For example, the parallel pattern of gene loss can be established in only 40 generations in *Tragopogon* (Buggs et al. 2012). Unlike angiosperms, the slow rate of chromosome loss in ferns may also indicate that many recombination-based deletion mechanisms may not play a significant role in ferns. A caveat to this is the rapid genomic downsizing of the model fern *Ceratopteris richardii*, which has fractionated rapidly and at a similar rate to many angiosperm taxa (Marchant et al. 2022). However, the general pattern in ferns seems to be similar to the Salmonids and *Xenopus*, in which diploidization is predominantly driven by pseudogenization (Berthelot et al. 2014; Lien et al. 2016; Session et al. 2016) and a slow rate of chromosomal change (Berthelot et al. 2014; Uno et al. 2013; Parey et al. 2022). Overall, our study provides additional evidence of more modest parallel and biased gene retention and loss following paleopolyploidy in monilophytes. Our observation of retention patterns might be consistent with slow rates of chromosome loss and the unique diploidization process in ferns.

## Conclusion

One of the most distinctive features of homosporous ferns is their high chromosome numbers relative to heterosporous plants (Klekowski and Baker 1966; Manton 1950). Using a comparative genomic framework, we provide confirmation of previous hypotheses that homosporous ferns likely have undergone a similar number of WGD as heterosporous plants, but fundamentally different mechanisms of diploidization. Our study provides structure for future work to reveal the potential processes driving the genomic differences among different lineages of vascular plants. It will be important in future work to include bryophytes in these analyses, as they are an entirely homosporous lineage with their own genome dynamics. Bryophytes have fewer rounds of WGD than angiosperms or pteridophytes (One Thousand Plant Transcriptomes Initiative 2019), but it is unclear how this has impacted their genome structure and chromosome number. Ongoing and future fern genome and transcriptome sequencing projects, especially on homosporous ferns with high chromosome numbers, will provide syntenic evidence to confirm putative ancient WGDs and better place these events in a phylogenetic context, advancing our understanding of the unique aspects of genome evolution in ferns.

## Methods

### DupPipe analyses of WGDs from transcriptomes of single species

For each transcriptome, we used the DupPipe pipeline to construct gene families and estimate the age distribution of gene duplications (Barker et al. 2010, 2008). We translated DNA sequences and identified reading frames by comparing the Genewise (Birney, Clamp, and Durbin 2004) alignment to the best-hit protein from a collection of proteins from 25 plant genomes from Phytozome (Goodstein et al. 2012). For all DupPipe runs, we used protein-guided DNA alignments to align our nucleic acid sequences while maintaining the reading frame. We estimated synonymous divergence (*Ks*) using PAML with the F3X4 model (Yang 2007) for each node in the gene family phylogenies. We identified peaks of gene duplication as evidence of ancient WGDs in histograms of the age distribution of gene duplications (*Ks* plots). We identified species with potential WGDs by comparing their paralog age distribution to a simulated null using a Kolmogorov–Smirnov goodness of fit test (Cui et al. 2006). We then used mixture modeling and manual curation to identify significant peaks consistent with a potential WGD and to estimate their median paralog *Ks* values. Significant peaks were identified using a likelihood ratio test in the boot.comp function of the mixtools R package(Benaglia et al. 2009).

### Estimating orthologous divergence

To place putative WGDs in relation to lineage divergence, we estimated the synonymous divergence of orthologs among species pairs that may share a WGD based on their phylogenetic position and evidence from the within-species *Ks* plots. We used the RBH Ortholog pipeline (Barker et al. 2010) to estimate the mean and median synonymous divergence of orthologs and compared those to the synonymous divergence of inferred paleopolyploid peaks. We identified orthologs as reciprocal best blast hits in pairs of transcriptomes. Using protein-guided DNA alignments, we estimated the pairwise synonymous (*Ks*) divergence for each pair of orthologs using PAML with the F3X4 model(Yang 2007). WGDs were interpreted to have occurred after lineage divergence if the median synonymous divergence of WGD paralogs was younger than the median synonymous divergence of orthologs. Similarly, if the synonymous divergence of WGD paralogs was older than that ortholog synonymous divergence, then we interpreted those WGDs as shared.

### Inference of ancient WGD in ferns

We explored the literature for studies that inferred WGD in ferns and lycophytes, and incorporated these findings into our analysis. We used four papers that included different sampling schemes, and therefore had varying estimates of WGD across pteridophytes. The One Thousand Plant Transcriptomes Initiative sequenced 1,124 green plants trancriptomes, including 59 fern and lycophyte species. Pelosi et al. (2022) used 247 transcriptomes to assemble 2,884 single-copy nuclear loci and infer WGD using *K*_s_ values. Huang et al. (2020) used 127 fern and lycophyte transcriptomes, and inferred genomes duplication using a linear regression of *K*_s_ values. Chen et al. (2022) inferred WGD in leptosporangiate ferns using gene tree-species tree reconciliation analyses as well as correcting for substitution rate. These studies detected many of the same WGD as our analyses did, but inferred four additional duplication events. As topology is known to vary between studies with different markers (e.g., (Hasebe et al. 1995; Schneider et al. 2004; Testo and Sundue 2016; PPG I 2016; McKibben, Finch, and Barker 2024), the placement of WGDs differed slightly across the studies used here. We used the K_s_ values to infer which WGDs matched across studies. If a *K*_s_ value was within 0.1 across all studies examined, we considered it to be the same WGD (see Fig. 1). One study (One Thousand Plant Transcriptomes Initiative 2019) inferred a WGD shared by all ferns at the base of the monilophyte phylogeny (WGD 1, see Supplementary Table 3). However, other studies have not inferred this WGD and there is no syntenic evidence for this putative ancient WGD. In order to remain conservative in our estimates, we did not count this in our total estimate of WGDs for ferns.

### Phylogenetic analysis of chromosome numbers

To estimate clade-wide rates of chromosome number change and to reconstruct ancestral chromosome numbers across the land plant phylogeny, we applied the likelihood-based phylogenetic method ChromEvol (Mayrose, Barker, and Otto 2010; Glick and Mayrose 2014) on a large combined data set of time-calibrated phylogenetic trees based on the *rbcL* gene (ribulose-1,5-bisphosphate carboxylase/oxygenase, large subunit) and chromosome numbers of the seed plants (angiosperms and gymnosperms) and ferns.

We constructed a genus-level phylogenetic data set for seed plants and ferns (Supplementary table 7). For each genus, both *rbcL* marker gene sequence data and chromosome counts available were represented. First, we retrieved *rbcL* sequences from NCBI GenBank (accessed Jul., 2011). The longest sequence (after disregarding Ns) among the sequences of the sampled taxa of a genus was used to represent the genus (randomly picking one sequence whenever there were ties). The taxon names were checked and corrected using the online Taxonomic Name Resolution Service version 5 (Boyle et al. 2013). Next, we downloaded chromosome numbers from CCDB version 1.46 (Rice et al. 2015); accessed June, 2020). The gametic chromosome number of a genus was summarized as the lower mode count of the gametic chromosome numbers of the sampled species; the gametic chromosome number of each sampled species was summarized as the lower mode count of the individual entries (including infraspecific entries) (Supplementary table 7). This way to summarize the chromosome count reduced probably erroneous chromosome counts. Also, the lower mode count of a species more likely reflected the common diploid cytotype of the species (Wood et al. 2009). Finally, we matched the *rbcL* sequence data and the summarized chromosome counts, retaining only the genera with both types of data available. The final data set contained the representative sequences and genus-level chromosome count summaries (aggregated from a total of 276,487 CCDB entries) of 2,531 genera (1,918 angiosperms, 50 gymnosperms, and 183 ferns) (Supplementary table 7). Additionally, the *rbcL* sequences of four lycophyte taxa were arbitrarily chosen to root the seed plant and fern phylogenies.

Using the *rbcL* sequences, we built two phylogenetic trees (seed plants and ferns) using PASTA version 1.7.8 (Mirarab, Nguyen, and Warnow 2014). PASTA was run using MAFFT version 7.149b (Katoh et al. 2002) as the multiple sequence aligner, OPAL version 2.1.3 (Wheeler and Kececioglu 2007) as the alignment merger, FastTree version 2.1.7 (Price, Dehal, and Arkin 2010) as the tree estimator (assuming GTR+G), and RAxML version 7.2.6 (Stamatakis 2014) to obtain the final tree after up to ten iterative refinements without improvement. The seed plant phylogeny contained 1,918 angiosperm genera and 50 gymnosperm genera and 15 arbitrarily selected fern and lycophyte outgroups (Supplementary table 7). The fern phylogeny contained 183 fern genera and 15 arbitrarily selected seed plants and lycophyte outgroups (Supplementary table 7). Both the *rbcL* phylogenetic trees were rooted with lycophytes, which were then pruned out from the phylogeny. Using PATHd8 version 1.0 (Britton et al. 2007), we calibrated the *rbcL* phylogenies. We took the fossil ages from four previous studies (Schneider et al. 2004; Smith, Beaulieu, and Donoghue 2010; Magallón et al. 2015; Testo and Sundue 2016), and applied the fossil ages as either fixed age or minimum age constraints in the PATHd8 analysis. We assigned as many of the fossil age constraints as possible to the internal nodes of the phylogenies, which corresponded to the most recent common ancestor of two taxa. Because we used genus-level phylogenetic data sets, fossil ages could not be assigned to the crown or stem group of the families or genera that were represented as a single tip on the phylogenies. Overall, we applied 86 fossil age constraints in the seed plant phylogeny and 17 fossil age constraints in the fern phylogeny (Supplementary table 8, Supplementary Figs. 4-5). For the ChromEvol analyses of each group ( ferns, gymnosperms, and angiosperms), we extracted the fern-only subtree from the fern time-calibrated phylogeny and the gymnosperm and angiosperm subtrees from the seed plant time-calibrated phylogeny.

ChromEvol models chromosome number change as a continuous-time Markov process along a phylogeny, in which observed chromosome numbers were assigned to each taxa. ChromEvol co-estimates the rates of ascending dysploidy (i.e., chromosome number gain or single increment), descending dysploidy (i.e., chromosome number loss or single decrement), duplication, and demi-duplication (i.e., multiplication by 1.5, e.g., the transition between tetraploids and hexaploids) via maximum likelihood. In this study, we employed the four ChromEvol models that assume that the rate of ascending dysploidy, the rate of descending dysploidy, the rate of duplication, and the rate of demi-duplication are constant throughout a given phylogeny. In the “CONST RATE” model, the rate of demi-duplication is fixed to zero while the rates of ascending dysploidy, descending dysploidy, and duplication are allowed to vary; in the “DEMI” model, the rate of duplication is set to equal to the rate of demi-duplication while the rates of gain and loss are allowed to vary; in the “DEMI EST” model, all of the four rate parameters are allowed to vary; and in the “NO DUPL” model, the rates of duplication and demi-duplication are both set to zero while the rates of gain and loss are allowed to vary (i.e., no polyploidy happens). We construe that there was no evidence for polyploidy in a data set if none of the three models allowing for duplication and/or demi-duplication was favored over the “NO DUPL” model.

We fitted the four ChromEvol models that assumed constant rates of ascending dysploidy, descending dysploidy, duplication, and demi-duplication. ChromEvol version 2 was run on the default optimization scheme, setting the minimum allowed chromosome number to one and the maximum allowed chromosome number to be twice the highest chromosome number observed in a given dataset. The upper limit was applied this way to ensure that the transition rate matrix is large enough to capture a plausible range of chromosome numbers (i.e., allowing doubling for the species with the highest chromosome number), while keeping the runs computationally feasible. For each clade, the best-fitting ChromEvol model was defined as the model having the lowest Akaike Information Criterion (AIC) score (Akaike 1974).

We estimated the clade-wide rates of polyploidy as the sum of the rate of duplication and the rate of demi-duplication and the clade-wide rates of dysploidy as the sum of the rate of descending dysploidy and the rate of ascending dysploidy. The rates of polyploidy and the rates of dysploidy were highly similar when estimated under the full model (i.e., “DEMI EST”) and the best-fitting model (which was not necessarily the full model). We used the results obtained under the best-fitting models (1) to assess ChromEvol model support across the clades and (2) to reconstruct ancestral chromosome numbers in the phylogenies of the clades. When comparing the ChromEvol rate estimates across the clades examined, we took the rates obtained under the full model to avoid biases in the rate estimates that could be introduced by forcing the rate of duplication and/or the rate of demi-duplication to be zero. The results of the ChromEvol analyses are presented in Supplemental Table 5.

We investigated how ancestral chromosome numbers changed over time in the angiosperms, gymnosperms, and ferns. We took the ancestral chromosome number inferred under the best ChromEvol model at each internal node (which had the highest posterior probability) and plotted the numbers over time (Fig. 3).

### Rates of protein evolution of meiosis genes in angiosperms and ferns

To estimate the rates of protein evolution, we select 352 meiosis-related genes based on GO terms under meiosis and their child genes. To avoid the impact of rate heterogeneity between angiosperms and ferns, we compare the rate of protein evolution for these meiosis genes to 500 randomly selected genes from the *Arabidopsis* genome. Meiosis-related genes and randomly selected genes are used as queries. Whole genome of 11 angiosperms (*Amaranthus hypochondriacus*, *Amborella trichopoda*, *Ananas comosus*, *Aquilegia coerulea*, *Arabidopsis thaliana*, *Carica papaya*, *Eucalyptus grandis*, *Mimulus guttatus*, *Oryza sativa*, *Populus trichocarpa*, and *Vitis vinifera*) were downloaded from Phytozome (Goodstein et al. 2012) used as the angiosperm database. Whole genome of *Azolla filiculoides* and *Salvinia cucullata* (F.-W. Li et al. 2018), and transcriptomes of eight ferns (*Asplenium formosae*, *Ceratopteris thalictroides*, *Dicksonia antarctica*, *Equisetum diffusum*, *Microlepia speluncae, Ophioglossum vulgatum*, *Osmunda javanica*, *Polypodium hesperium*) were used as the database for ferns. *Physcomitrella patens* are used as an outgroup, and *A. thaliana* and *P. patens* were outgroups in fern analyses. Both queries were used to blast against the angiosperm and fern databases using protein BLAST (blastp) with E-value of 1e-20 as a threshold. The hits from each protein blast search were used for gene family clustering with Orthofinder 2.3.7 (Emms and Kelly 2015). Gene families with the presence of at least one *P. patens* sequence and one *A. thaliana* sequence were selected. PASTA (Mirarab, Nguyen, and Warnow 2014) was used to construct gene family phylogenies. Gene trees are re-rooted by the *Physcomitrella patens* and tips of outgroups are dropped in the gene trees after re-rooting. The root-to-tip distance is extracted with the ‘distances’ function in ape (Paradis and Schliep 2019). For each gene tree, we estimate the mean, minimum, and maximum root-to-tip distance for each species. To compare the rate of protein evolution, we use the Mann-Whitney *U* test to compare the distribution of root-to-tip distance between meiosis and randomly selected genes within angiosperms and ferns. To visualize these distributions, we also plot the distributions of each comparison of angiosperms and ferns with ggplot in R.

### Gene Ontology (GO) annotations and paralog retention and lost patterns

To obtain gene ontology (GO) annotations, we followed the methodology of the GOgetter pipeline (Sessa et al. 2023). GO annotations of all fern transcriptomes were obtained through discontiguous MegaBlast searches against annotated *Arabidopsis thaliana* transcripts from TAIR(Swarbreck et al. 2007) to find the best hit with a length of at least 100 bp and an e-value of at least 0.01. We evaluated the overall differences between the GO composition of transcriptomes and WGD paralogs by principal component analysis (PCA) using the rda function in vegan (Dixon 2003). We tested if the GO category composition was different between transcriptome and WGD paralogs using a permutation test in the vegan envfit function. Ellipses representing the 95% confidence interval of standard deviation of point scores were drawn on the PCA plot using the ordiellipse function in vegan. We further tested for differences among GO annotations using chi-square tests. When chi-square tests were significant (*p* < 0.05), GO categories with residuals >|2| were implicated as major contributors to the significant chi-square statistic. A category with residual >2 indicates significant over-retention of this category following WGD, whereas residual <–2 indicates significant under-retention (Barker et al. 2008). Using this statistical framework, we tested for significant differences between the overall transcriptome and paralogs from the all paleopolyploid species.

## Acknowledgments

We thank Y. Li for providing the illustrations of ferns. We also thank C. Roman-Palacios and members of the Barker lab for the discussion. S.P.K is funded by a National Science Foundation Postdoctoral Research Fellowship.

## Author contributions

Z. L., S.H.Z., and M.S.B. designed research; Z. L., S.H.Z., S.P.K, and M.S.B. performed research and analyzed data; Z. L., S.P.K, and M.S.B. wrote the paper.

## Supplemental Tables

**Supplementary Table 1 Summary of Ks distributions of duplicate gene pairs for each species.** Species are organized in alphabetical order. Taxa and median Ks for each histogram of the age distribution of gene duplications (Ks plots available at https://gitlab.com/barker-lab/fern-wgds-and-chromosomes), and p-values of the two-sided K-S goodness of fit test is provided for each taxon. The number of inferred WGD in the ancestry of each species is also reported. The phylogenetic placement of each inferred ancient WGD is provided in Figure 1 and Supplementary Fig. 1.

**Supplementary Table 2 Summary of the synonymous ortholog divergence analyses.** The mean, median, and standard deviation of synonymous ortholog divergence (Ks) for each inferred ancient WGD is reported. These statistics are based on the number of ortholog pairs identified for each species pair comparison. The sampling information with taxon code and WGD code is also reported.

**Supplementary Table 3 Ancient WGDs inferred in the phylogeny of ferns.** The phylogenetic placements and support from different studies are provided. The WGD code matches with WGD placements on Supplementary Fig. 1

**Supplementary Table 4 Rates of ancient genome duplication in ferns and other vascular plants.** The rates of ancient WGD/Ma were estimated using the number of ancient genome duplications inferred by the genomic analyses divided by the minimum crown group age for each vascular plant lineage (Fig. 2a). The rates of ancient WGD/Ma estimated by ChromEvol are provided for comparison.

**Supplementary Table 5 Results of the ChromEvol analyses of angiosperm orders, gymnosperms, and monilophytes.** The best-fitting model (out of “NO DUPL”, “CONST RATE”, “DEMI”, and “DEMI EST”; see Methods section for a description of the models) was inferred based on the lowest AIC score. The rates of chromosomal evolution (in events per a million years) estimated under the best-fitting model and the full four-parameter model (“DEMI EST”) are provided. The rate of dysploidy was defined as the sum of the rate of chromosome loss (−1) and the rate of gain (+1). The rate of polyploidy was defined as the sum of the rate of duplication (2x) and the rate of demi-duplication (1.5x). Additionally, for each clade, the ancestral chromosome number inferred at the root of the clade’s phylogenetic tree under the best-fitting model or the four-parameter model (“DEMI EST”) are indicated.

**Supplementary Table 6 Sample size and summary statistic for rate of protein evolution of meiosis-related genes in angiosperms and ferns.** The distribution of root-to-tip distance between meiosis and background genes in angiosperms and ferns was estimated (see methods). The root-to-tip distances were calculated using maximum, mean, and minimum, with and without excluding outliers. The comparisons of the mean of meiosis and random genes by the two-sample Mann-Whitney U test are provided. The mean, median, standard deviation, U, Z, and p-value are provided for each root-to-tip distance estimate.

**Supplemental Table 7 rbcL accession numbers and gametic count for the seed plant and fern ChromEvol analyses.** Genera in the phylogenetic analysis of the seed plants and ferns were used. The taxonomic order (as per APG IV or PPG I classification), family (as per TROPICOS classification), and representative taxon of each genus examined are shown alongside the GenBank accession of the *rbcL* sequence of the representative taxon. Also indicated are the gametic chromosome number of each genus (summarized as the lower mode of species-level gametic chromosome numbers, which were also summarized using the lower mode) and the number of CCDB entries summarized for the genus. Taxonomic name resolution was attempted using TRNS (Taxonomic Name Resolution Services V5.0), and synonymous names were recorded when applicable. The genera used as the outgroup taxa were noted.

**Supplemental Table 8 Fossil records used in PATHd8 for the seed plant and fern ChromEvol analyses.** The fossil ages of the crown or stem group of various clades were taken from Schneider et al. (2004) *Nature*, Magallón et al. (2015) *New Phytol*, and Smith et al. (2010) *PNAS,* Testo and Sundue (2016) *Mol Phylogenet Evol*. In a PATHd8 analysis, the fossil ages were employed as a fixed age constraint or minimum age constraint on the internal node representing the most recent common ancestor of a focal clade, which was defined by two tip taxa in the phylogeny. See Supplementary Fig. 3-5 for the placement of the fossil age constraints.

## Supplementary Figures

**Supplementary Fig. 1 Enumerated ancient WGDs inferred in the phylogeny of ferns.** Each of the numbered circles corresponds to the WGD shown in Figure 1. These numbers are used to refer to each WGD in Supplementary Tables 1 and 3. Dotted circles show where previous authors placed WGD in their trees, but the Ks values of those WGD are very close to what we inferred, so we believe they are the same duplication (dotted lines connect the two). This phylogeny was modified from Testo and Sundue, 2016.

**Supplementary Fig. 2 GO annotations of whole transcriptomes and genes retained from ancient WGDs.** Each column represents the annotated GO categories of pooled whole transcriptomes or genes retained in duplicate following ancient WGD from each analyzed species. Colors of the heatmap represent the percent of the transcriptome represented by a particular GO category. The overall ranking of GO category rows was determined by the ranking of GO annotations among the total pooled transcriptomes. Hierarchical clustering was used to organize the heatmap columns.

**Supplementary Fig. 3 Pattern of gene retention and loss following ancient WGDs.** Each column represents the annotated GO categories of paralogs retained following ancient WGDs from each analyzed species. The order of analyzed species (20 hexapods and one outgroup) are based on hierarchical clustering. The overall ranking of GO category rows was determined by the ranking of GO annotations among ancient WGD 10 in *Osmundastrum cinnamomeum*. Colored boxes indicate GO categories among putative WGD paralogs that were significantly over- (red) or under-retained (blue) relative to the pooled whole transcriptomes, as determined by residuals from chi-square tests. GO categories with gray boxes were not present among WGD paralogs in significantly different numbers relative to their frequency in the pooled whole transcriptomes.

**Supplementary Fig. 4 *rbcl* phylogeny of seed plants.** We applied 86 fossil age constraints in the seed plant phylogeny and the placements of the fossil age constraints are shown on the phylogeny.

**Supplementary Fig. 5 *rbcl* phylogeny of ferns.** We applied 17 fossil age constraints in the fern phylogeny and the placements of the fossil age constraints are shown on the phylogeny.

**Supplementary Fig. 6 Patterns of inferred ancestral chromosome number and posterior complement values in angiosperms, gymnosperms, and ferns.** The right-hand x-axis represents the root-to-internal node distance on the phylogenies analyzed using ChromEvol. The left-hand x-axis shows the respective posterior complement (credible intervale) values for each inferred chromosome number. The y-axis shows the ancestral chromosome numbers taken from the ChromEvol results. The dotted line shows the median chromosome number for each group. The rate of polyploidy, ascending and descending dysploidy are presented in the figure legends. **a**. 1,918 data points show the pattern in angiosperms, median *n*=10. **b**. 51 data points show the pattern in gymnosperms, median *n*=11. **c**. 199 data points show the pattern in ferns, median *n*=33. The decrease in the posterior complement through time for each group suggests that other factors might be contributing to extant chromosome numbers.

## Data Availability

All files for the analyses presented in this paper can be found at https://gitlab.com/barker-lab/fern-wgds-and-chromosomes.

## Input and output files of the chromosome number analyses

The ChromEvol analyses were conducted in three stages: 1) PASTA, 2) PATHd8, and 3) ChromEvol. First, the maximum-likelihood phylogenies of the ferns and seed plants (angiosperms and gymnosperms) were inferred separately using PASTA. Then, these two ML phylogenies were dated separately using PATHd8. Finally, ChromEvol was run separately on the fern phylogeny, gymnosperm phylogeny (which was extracted from the dated seed plant phylogeny), and the angiosperm order phylogenies (which were extracted from the dated seed plant phylogeny).

## Rates of protein evolution of meiosis genes in angiosperms and ferns

To compare the rate of protein evolution, we compared meiosis and randomly selected genes within angiosperms and ferns. We selected 352 meiosis-related genes based on GO terms under meiosis and their child gene. We randomly selected 500 genes from the *Arabidopsis* genome. Provided are the trees for testing the rates of molecular evolution.

## Ks distributions of duplicate gene pairs

140 Histograms of the age distribution of gene duplications (Ks plots) range from Ks 0 to 2, and (plotted separately) Ks 0 to 5. The phylogenetic placement of each inferred ancient WGD is provided in Figure 1 and Supplementary Fig. 1.

