## Supplementary figures and images for "Ancient polyploidy and low rate of chromosome loss explain the high chromosome numbers of homosporous ferns"

### SI_Figure_1_fern-tree-wgd-numbers.pdf

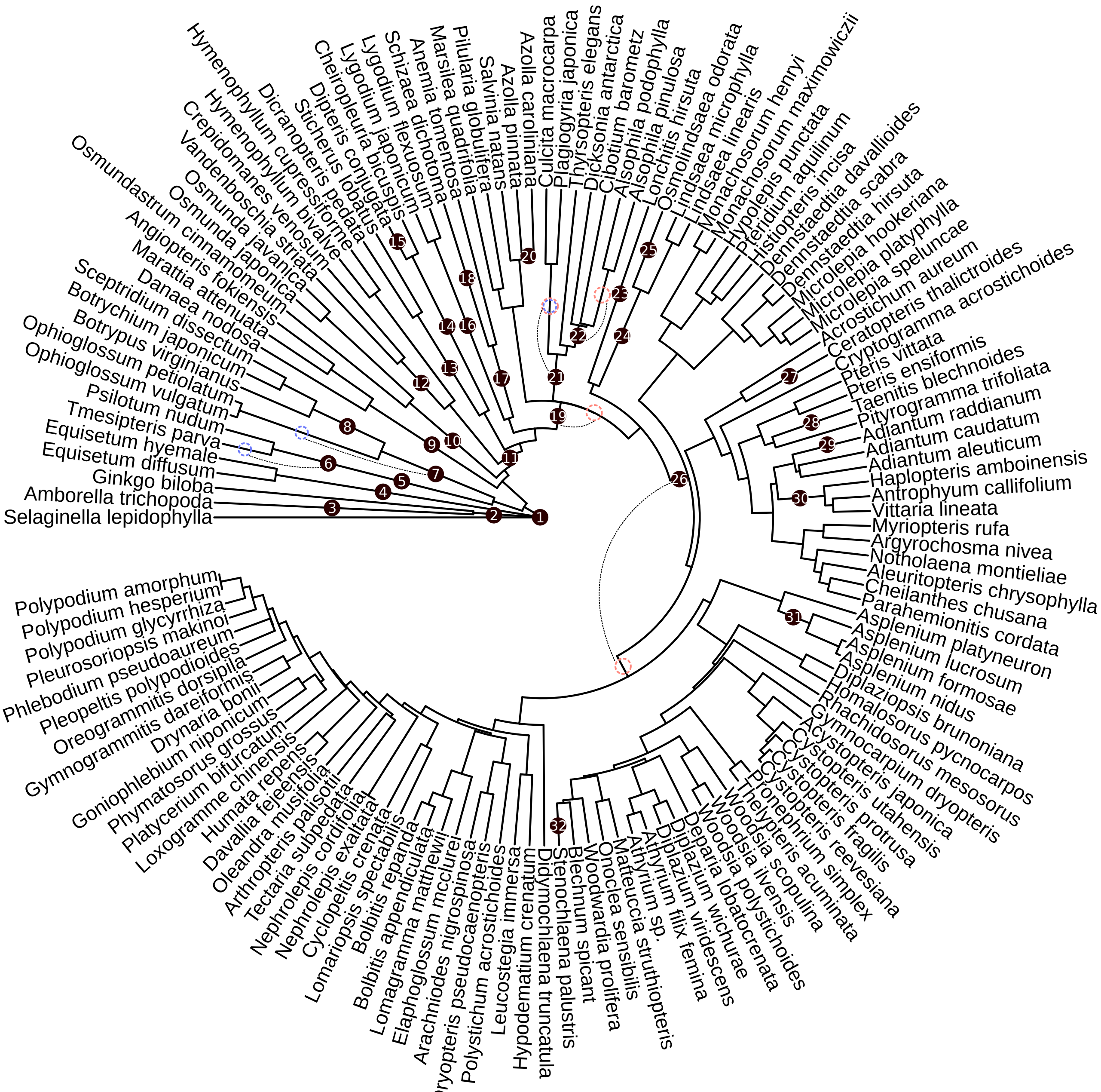

### SI_Figure_2_freq_26_wgd_in_r.png

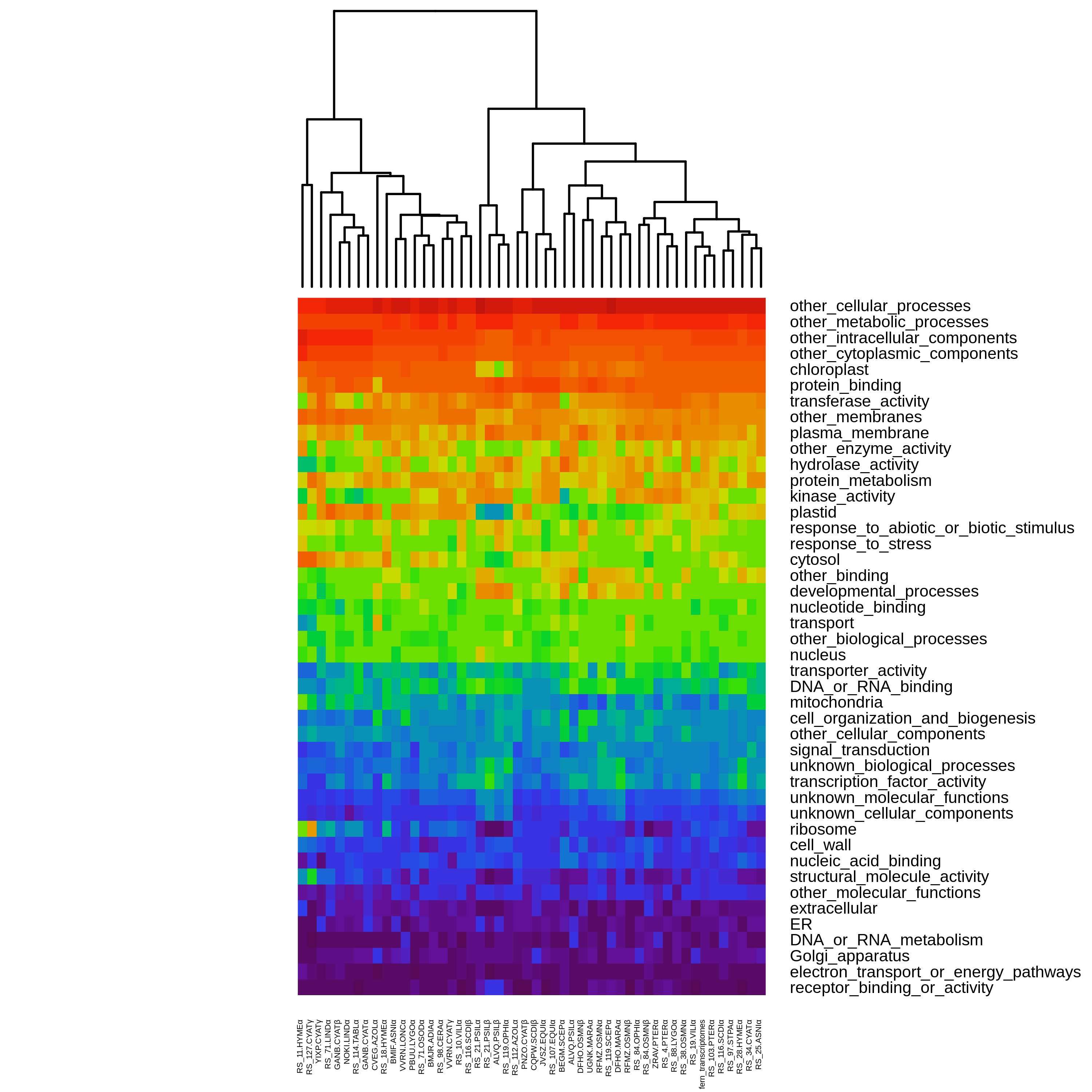

### SI_Figure_3_freq_26_wgd_in_r.png

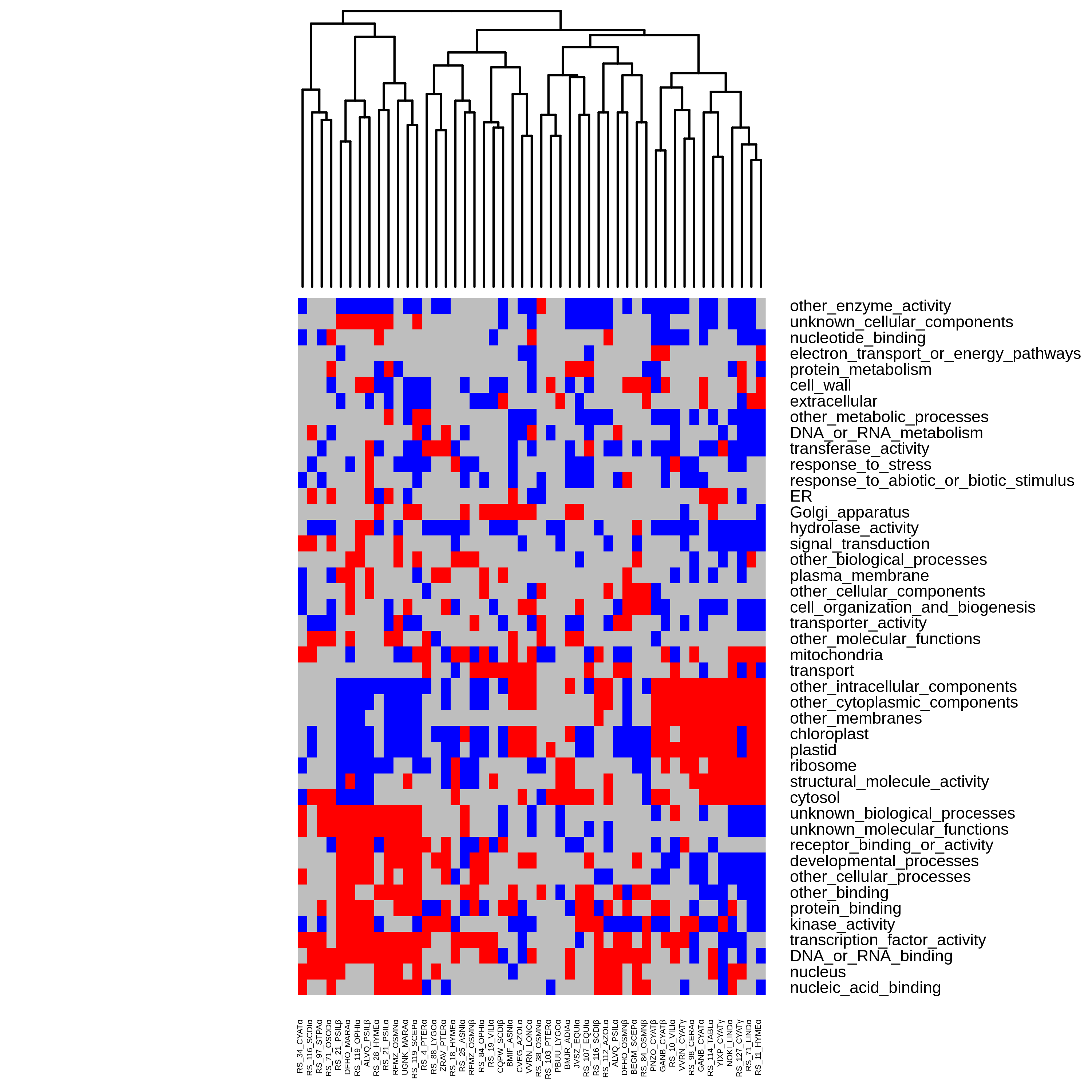

### SI_Figure_5_fern_timetree.pdf

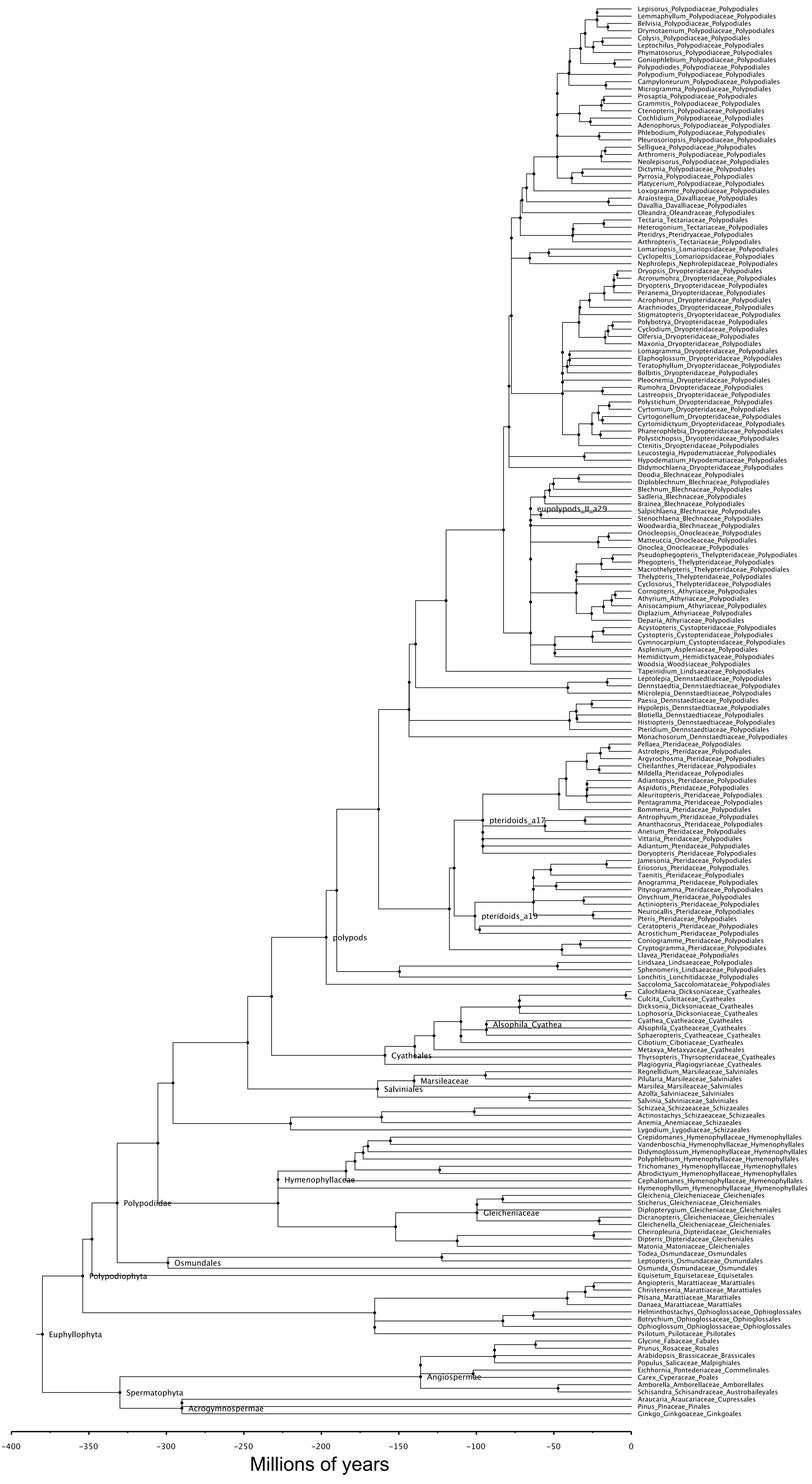

### SI_Figure_6_chromosome_number_posterior_complement.pdf

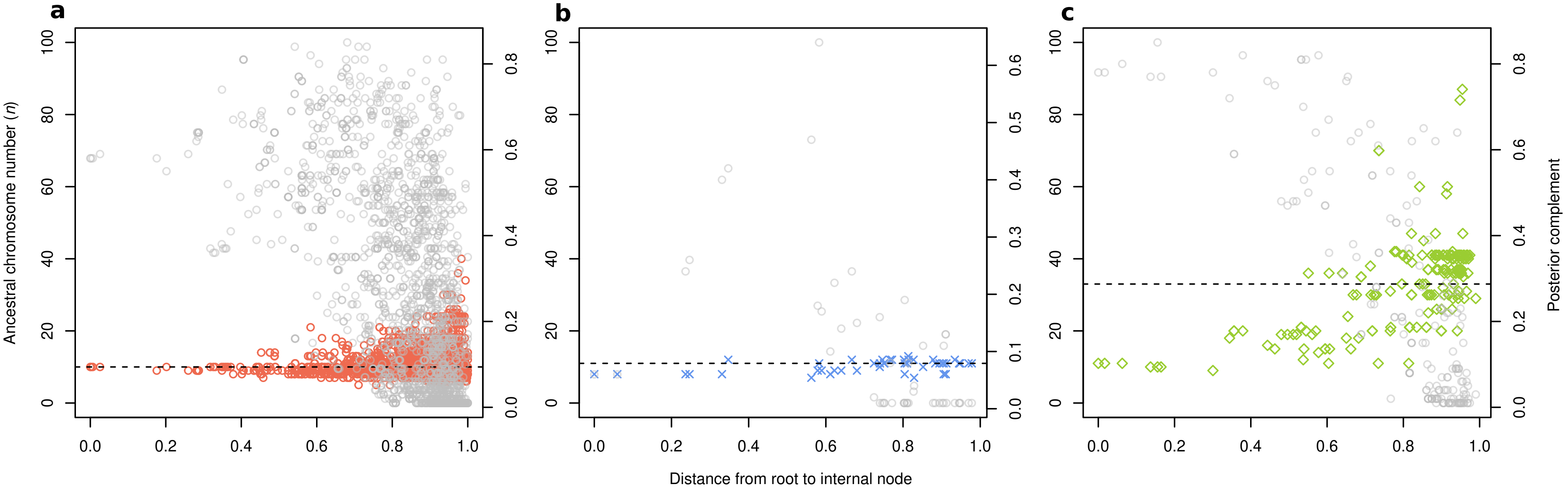
